# Topology and cooperative stability: the two master regulators of protein half-life in the cell

**DOI:** 10.1101/181925

**Authors:** Saurav Mallik, Sudip Kundu

## Abstract

In a quest for finding additional structural constraints, apart from disordered segments, regulating protein half-life in the cell (and during evolution), here we recognize and assess the influence of native topology of biological proteins and their sequestration into multimeric complexes. Native topology acts as a molecular marker of protein’s mechanical resistance and consequently captures their half-life variations on genome-scale, irrespective of the enormous sequence, structural and functional diversity of the proteins. Cooperative stability (slower degradation upon sequestration into complexes) is a master regulator of oligomeric protein half-life that involves at least three mechanisms. (i) Association with multiple complexes results longer protein half-life; (ii) hierarchy of complex self-assembly involves short-living proteins binding late in the assembly order and (iii) binding with larger buried surface area leads to slower subunit dissociation and thereby longer half-life. Altered half-lives of paralog proteins refer to their structural divergence and oligomerization with non-identical set of complexes.

---

Cellular proteins are regularly degraded and replaced with newly synthesized copies, minimizing the accumulation of toxic damage and ensuring a functional proteome. An elegant balance between translation and degradation rates thus maintains protein concentration within the cell, assigning each protein a specific half-life^1–4^. A protein’s life starts as its messenger RNA blueprint is translated into a chain of amino acid building blocks. This chain generally folds itself into a 3D molecule that then takes on functions such as enzymatic activity, binding specific ligands, helping to create cellular structures, assembling into macromolecular machines and transporting other proteins. Protein’s life ends as a degradation machinery, such as the ubiquitin-proteasome system (UPS) in eukaryotes, proteolyzes it into multiple fragments^4,5^. The UPS includes two major enzymes. One is ubiquitin, that stochastically festoons substrate proteins with a molecular marker for degradation (a polyubiquitin tag). The other is proteasome, that (i) recognizes its substrates based on this tag, (ii) engages with an intrinsically disordered region (IDR) of the substrate, (iii) mechanically unfolds the protein by pulling the polypeptide chain from the engaged IDR into a degradation channel^6^ where (iv) an ATP-driven proteolysis occurs^5–7^ (Fig. 1a).

**Figure 1.**
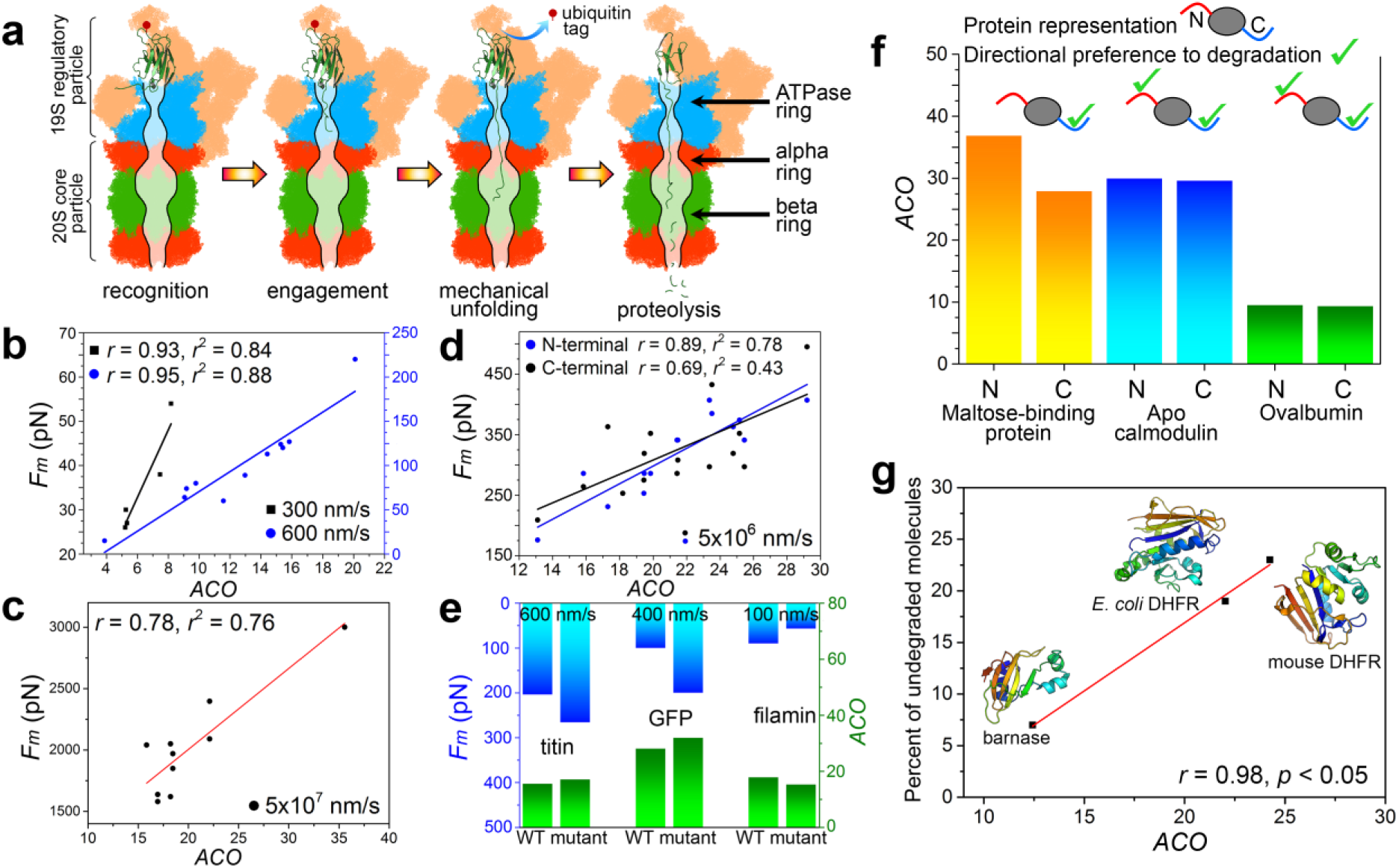
Native topology acts as a marker of protein’s mechanical resistance. (**a**) A schematic representation of proteasome function. (**b**) The peak unfolding forces estimated for pulling the termini of 11 globular proteins at 600 nm/s and of 5 proteins at 300 nm/s speeds (G1 set), in Atomic Force Microscopy experiments, are plotted against their native *ACO*. (**c**) The peak unfolding forces estimated in all-atom computer simulation to unwind 11 proteins from N-terminal at 5 × 10^7^ nm/s speed are plotted against their native *ACO* (G2A set). Solid lines indicate linear regressions. (**d**) The peak unfolding forces estimated in all-atom computer simulation to unwind 16 proteins from N- and C-terminal separately at 5 × 10^6^ nm/s speeds are plotted against their native *ACO* (G2B set). (**e**) For titin, green fluorescent protein and filamin, elevation/demotion of mechanical resistance upon point-mutations is associated with alike changes of *ACO*. The three proteins are unfolded at three different speeds, suggesting this pattern is irrespective of chain-pulling speeds. (**f**) Proteasome prefers to unwind globular proteins by using the terminal as initiation site that requires minimum peak unwinding force. For three such experimentally verified cases, where the two termini are located at two distinct structured domains, *ACO* associated with the two domains are plotted, along with highlighting the directional preference of proteasome. (**g**) The percent of undegraded molecules of barnase and dihydrofolate reductase (DHFR, from *Escherichia coli* and mouse) after 200 minutes of incubation with the proteasome are plotted against their *ACO*.

Experimental measurement of protein half-life in different organisms show a wide range of variation from minutes to days^1–3^, providing a platform based on which multiple biological questions can be addressed. Some studies have shown altered protein half-lives leading to abnormal development^8^, neurodegenerative diseases and cancer^9^. Accumulation of toxic damage in long-lived proteins is identified as a major inducer of ageing^10^. Other studies have looked for the factors that affect protein half-life in the cell^11–19^. Over the years, multiple factors have been identified—some tested only for specific proteins, some tested at genome-scale—to affect protein half-life in the cell.

The proteolytic site of proteasome is accessible only through a narrow degradation channel (10–15Å width, ∼70Å length), through which only unstructured polypeptides can penetrate^4,5^. Consequently, on a genome-scale, proteins featuring long intrinsically disordered regions (IDRs) are more susceptible to degradation and they exhibit short half-lives^12^. Shorter half-life is also observed for proteins featuring IDRs with amino acid compositions permitting high-affinity proteasomal engagement^11^. To degrade globular proteins, the ATPase molecular motor of proteasome first sequentially unfolds them by pulling their polypeptide chain from the engaged IDR into the degradation channel^4,13^. This mechanical unfolding is resisted by the native molecular contacts stabilizing the globule^14^ and only for a handful of proteins, it is shown that stronger resistance leads to slower degradation rates^15,16^. Protection from degradation is also achieved when proteins sequestrate into multicomponent complexes^17–19^. This effect is formally known as cooperative stability^20^, but neither its molecular basis is clearly understood, nor its impact on protein half-life is tested on a genomic scale.

Here, we exploit the experimental genome-scale half-life data of yeast proteins, wide-ranging information about their structural fold and 3D geometry, along with extensive biochemical characterization of the complexes they assemble into to develop a theory demonstrating how a wide spectrum of structural constraints of biological macromolecules regulates protein half-life in the cell. We begin by finding that native topology of monomeric globular proteins acts as a molecular marker of their mechanical resistance, and thus, affects half-life on a genomic-scale. For oligomeric proteins, the influence of topology is superseded by that of cooperative stability, that affects half-life in at least three mechanisms, (i) association with multiple complexes leads to longer half-lives of subunit proteins, (ii) hierarchy of complex self-assembly involves short-living proteins binding late in the assembly order and (iii) for small complexes, larger buried surface area, that generally reflects strong association and weak dissociation constants, generally leads to longer half-lives. Finally, we confirm that diversification of native topology and promiscuous oligomerization are further exploited to alter protein half-life during evolution. Our work not only evaluates the independent and combined impacts of different structural constraints to regulate protein half-life, and places them into genomic context, but further deepens our understanding of the designing principles of biological macromolecules.

## RESULTS

### Prevalence of long-range contacts of globular proteins contribute to stronger mechanical resistance and thereby longer half-life

Mechanical unfolding is a crucial step of globular protein degradation^4,13^ and this phenomenon has received a great deal of scientific focus in the past decade, encouraging multiple experimental and simulation studies attempting to understand the molecular origin of protein’s mechanical resistance (Data S1). An interesting comparison of ubiquitin and protein L (similar fold class) showed equivalent unfolding patterns at all chain pulling speeds, but the former having higher native long-range contacts (non-covalent contacts between residues far separated in primary chain) required higher peak unfolding force^21^. Starting from this point, we ask to what extent the prevalence of long-range native contacts of globular proteins (quantified as absolute contact order (*ACO*), the average primary chain separation of atomic contacts) affect their mechanical resistance. We perform three analyses. First, we estimate the correlation between native state *ACO* and the peak force required for mechanical unfolding (*Fm*) for two groups of proteins (Online Methods). The first group (G1) includes 16 proteins unfolded in Atomic Force Microscopy experiments, by pulling the polypeptide chains at 600 nm/s (11 proteins) and 300 nm/s (5 proteins) speeds. The second group (G2) includes 27 proteins unfolded in all-atom computer simulations. Eleven proteins (G2A) were pulled from the N-terminal at 5 ×10^7^ nm/s speed (C-terminal fixed). Sixteen proteins (G2B) were pulled from N- and C-terminal separately at 5 ×10^6^ nm/s speed, keeping the other terminal free, thus allowing the substrate to rotate and adopt a less obstructive orientation for unfolding (as happens during degradation). Between *ACO* and *Fm*, we obtain surprisingly strong 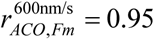 and 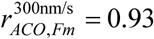 correlations in G1, 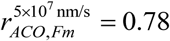 in G2A and 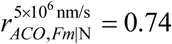, and 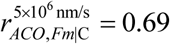 in G2B for N- and C-terminal pulling respectively (**Fig. 1b-d**). Second, for three globular proteins with experimental data depicting alterations of mechanical resistance upon point-mutations (Online Methods), we confirm that an elevation/demotion of mechanical resistance is perpetually associated with alike changes of *ACO* (**Fig. 1e,** Data S1). Third, for some proteins it was demonstrated that their mechanical anisotropy (pulling from different termini requires different peak unfolding forces) determines the directional bias of degradation (the terminus that is intrinsically disordered / easier to mechanically unfold is preferred by the proteasome to initiate degradation)^22^. For three such cases (maltose-binding protein, apo-calmodulin and ovalbumin), where the two termini are located at two distinct structured domains, we make two crucial observations. (i) Proteasome prefers unwinding maltose binding protein from the C-terminal domain, that has lower *ACO* (and requires weaker unwinding force) compared to that of N-terminal domain. (ii) Proteasome has no directional preference to unwind apo-calmodulin and ovalbumin, and both of their N- and C-terminal domains exhibit nearly identical *ACO* (**Fig. 1f**). These three sets of analyses provide a statistical proof-of-concept that *ACO* acts as a molecular marker of protein’s mechanical resistance, in a manner that higher *ACO* dictates higher mechanical resistance.

During an interesting experiment of titin degradation by ClpXP (bacterial/mitochondrial homolog of proteasome) Kenniston et al.^15^ observed that folded titin molecules are processed at much slower rates (150 molecules/min) than unfolded ones (600 molecules/min). They concluded that proteasomal degradation being a stochastic process, each substrate has a fixed probability of denaturation during each enzymatic cycle. For substrates with stronger mechanical resistance (such as folded titin, compared to unfolded ones), this probability would be lower and denaturing most of the molecules in the population would require many ATP cycles^15^. Since higher *ACO* prompts higher mechanical resistance, for two proteins subjected to proteasomal degradation for the same time span, larger fraction of undegraded molecules is expected for the one with higher *ACO*. This notion is supported by the outcome of an experiment subjecting dihydrofolate reductase (from *Escherichia coli* and mouse) and ribonuclease barnase proteins to proteasomal degradation^13^. After 200 minutes of incubation, the percent of undegraded molecules of the three proteins exhibit a surprising −0.98 correlation with their *ACO* values (**Fig. 1g**). These results encourage us to ask whether and how native topology influences protein half-lives in the cell. We start with 52 X-ray crystallographic structures (≤3 Å resolution) of annotated yeast monomeric proteins (sequence coverage of crystal structure *sc* = 100%) and obtain a surprising 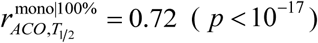 correlation between *ACO* and log *T*_1 /2_ (**Fig. 2a**). For the 158 oligomeric protein structures as well, collected under the same criterion, we find a statistically significant, albeit much weaker correlation (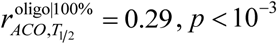, **Fig. 2b**). Even if we include crystal structures of 30 monomeric and 71 oligomeric proteins with missing coordinates (signify flexible or disordered regions and crystal artifacts, *sc* ≥ 75% is taken), significant correlations are obtained (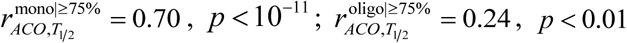, **Fig. 2c-d**).

**Figure 2.**
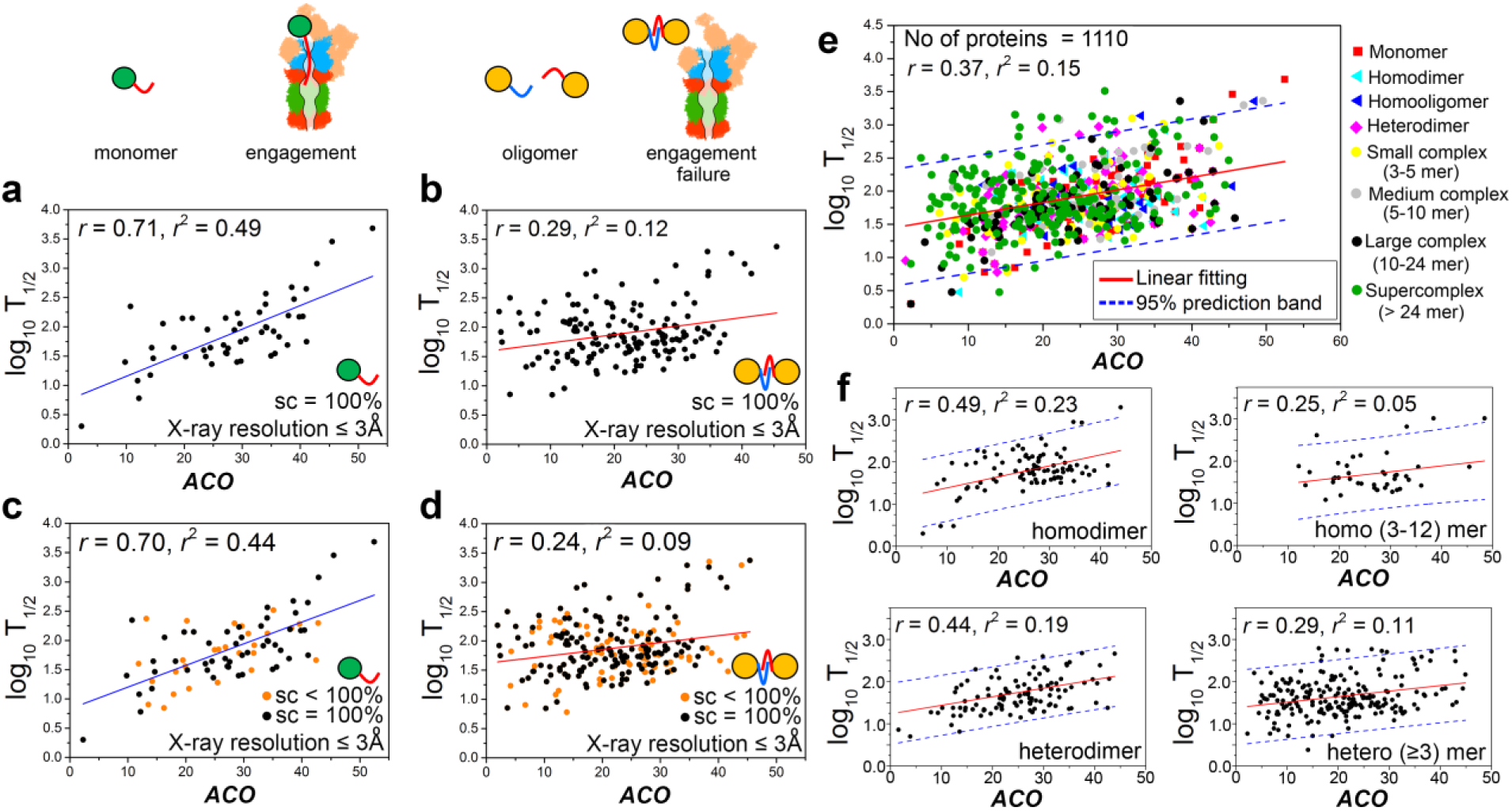
Native topology captures half-life variations of globular proteins on a genomic scale. (**a**) For monomeric and (**b**) oligomeric proteins with crystal structures covering entire protein lengths, logarithms of half-life values are plotted against native state *ACO*. Solid lines signify linear regression. (**c**) For monomeric and (**d**) oligomeric proteins with crystal structures covering ≥ 75% of protein lengths, logarithms of half-life values are plotted against native state *ACO*. (**e**) For crystal and modeled structures of 1110 yeast proteins, their logarithmic half-lives are plotted against their native state *ACO*, followed by a linear regression. (**f**) Four plots depict that for larger complexes, the correlation between logarithmic half-life and *ACO* drops. The top panels include the plots for homodimers and homooligomers, the bottom panels include plots for heterodimers and heterooligomers.

Even after including all the ≤3 Å resolution structures with *sc* ≥ 75%, we are left with structures of only 311 proteins, which although depicts a significant correlation (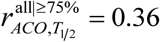, *p* < 10^-13^), but is inadequate to infer a proteome-wide tendency (*T*_1/ 2_ known for 3274 proteins). Hence, we extend our structure dataset by including additional 799 modeled structures (*sc* ≥ 75%) generated with reliable accuracy of fold assignment (Online Methods). For this set of total 1110 crystallographic and modeled structures, we obtain a striking 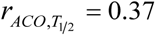 correlation (*p* < 10^-37^) between log *T*__1 2__ and *ACO*, demonstrating a proteome-wide tendency of native topology regulating protein half-lives (**Fig. 2e**).

Notably, 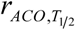 is stronger for monomers (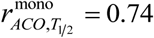, *p* < 10^-18^), compared to both homo- (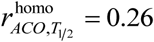, *p* < 0.01) and heteromers (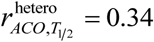, *p* < 10^-13^), for any *sc* ≥ 75%. Molecular basis of this weak correlation probably refers to at least two factors. First, the cooperative stability^20^ of oligomeric proteins (escaping proteasomal degradation in complexed state^17–19^) is generally independent of, and often overpowers, the effect of *ACO*. Degradation of β-casein is an interesting example of this trend. Intrinsically disordered C-terminal domains of two β-casein molecules dock together to form a homodimer, forcing the proteasome to initiate degradation exclusively from the globular N-terminus^22^. Second, proteins associated with larger complexes (multiple subunits) are generally more flexible and experience higher structural rearrangements upon oligomerization^23^. Oligomeric proteins are observed to degrade much faster at their monomeric states^17–19^, the *ACO* of which is not equal to the *ACO* we estimate from crystal structure data of yeast complexes. This may also weaken 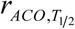 correlations in oligomeric proteins. This notion is supported by the gradual reduction of 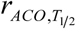 for larger homo- and heteromeric complexes (**Fig. 2f**).

Taken together, these data show that native topology acts as a master regulator of globular protein half-life, with indications that cooperative stability has some strong influence as well.

### Promiscuity of oligomerization results longer half-lives

To assess the impact of cooperative stability on oligomeric protein half-life, first we develop a proteome-scale database of yeast macromolecular complexes. Starting from earlier published databases^24,25^, we continue a protein-by-protein manual curation of available experimental data (Online Methods), yielding a massive database of 805 heteromeric and 80 homomeric yeast complexes (Data S2). This database includes 2487 annotated yeast proteins. Complex subunits are classified into two classes, central (functional subunits of a matured complex, if different isoforms of the complex exist^24,25^, they are present in most isoforms) and attached (temporary attached particles such as assembly cofactors, chaperones and subunits present in some of the isoforms).

First, we test whether sequestration into multicomponent complexes has any measurable impact on protein half-life, apart from that of their *ACO*. We classify mono-and oligomeric proteins into distinct groups based on their *ACO*, and compare the respective half-life distributions. For similar ranges of *ACO*, oligomeric proteins exhibit significantly longer half-lives than monomeric proteins (**Fig. 3a**), demonstrating cooperative stability is another master regulator of protein half-life across the genome^20^.

**Figure 3.**
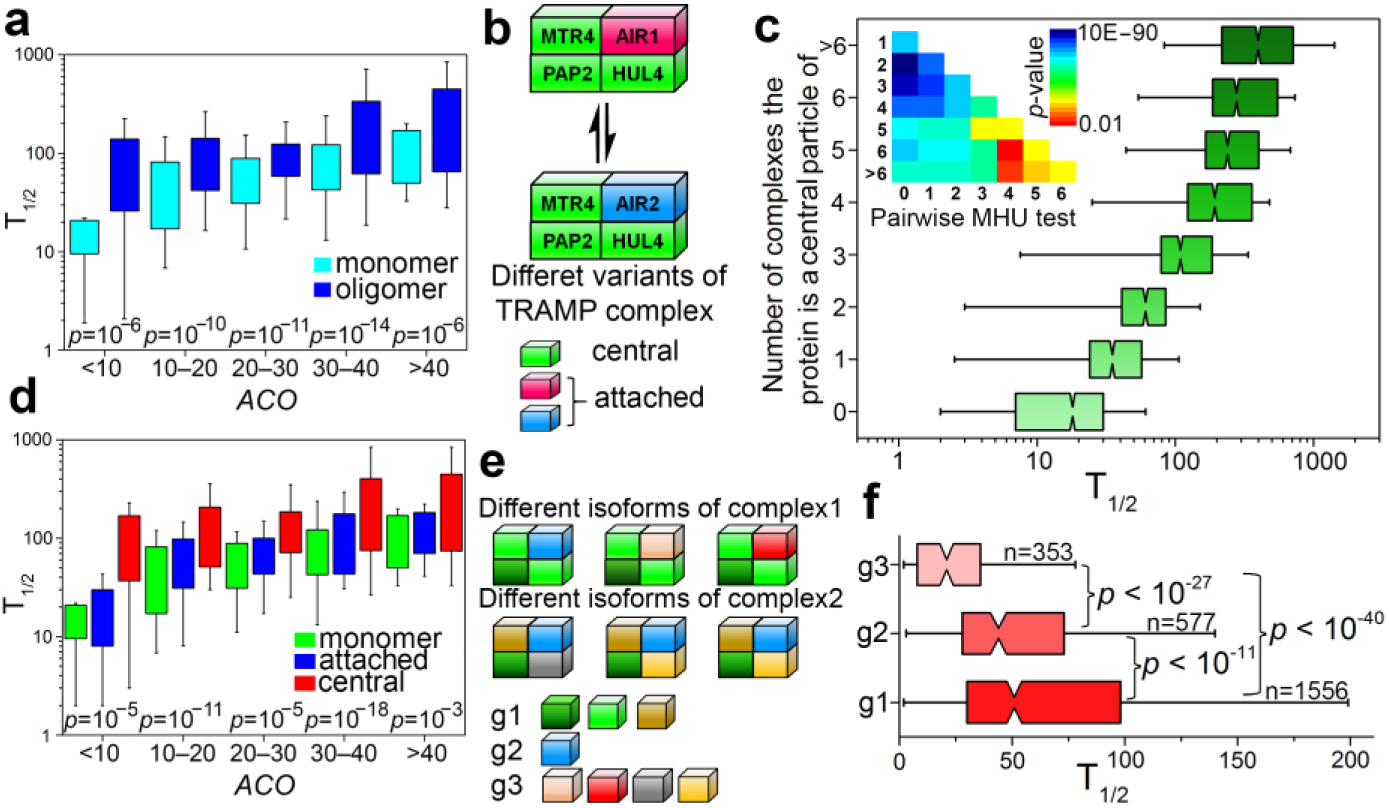
Cooperative stability is a master regulator of oligomeric protein half-life. (**a**) At similar ranges of native state *ACO*, half-life distributions of mono-and oligomeric proteins are compared using pairwise Mann-Whitney U-tests. (**b**) An example of different isoforms of TRAMP complex and definitions of central and attached subunits. (**c**) Comparing half-life distributions of proteins associated with different number of complexes (as central subunits) using pairwise Mann-Whitney U-tests. (**d**) At similar ranges of native state *ACO*, half-life distributions of monomeric, central and attached proteins are compared using pairwise Mann-Whitney U-tests. (**e**) A schematic representation of classifying oligomeric proteins into g1 (participate in ≥1 complexes as central particles only), g2 (contribute to ≥1 complexes as central and to ≥1 complexes as attached particles) and g3 (participate in ≥1 complexes as attached particles only) groups. (**f**) Comparing half-life distributions of proteins across g1, g2 and g3 using pairwise Mann-Whitney U-tests.

How does cooperative stability relate to complex size and involvement of proteins in different complexes? Earlier we observed weaker 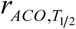 for complexes with multiple subunits. But surprisingly, participation in larger complexes is not associated with longer half-lives (Kruskal-Wallis (KW) test *p* > 0.05, which extends Mann-Whitney-U test to ≥2 groups). Rather promiscuity of oligomerization appears to be a strong modulator of cooperative stability, in a matter that involvement in higher number of complexes as central particles is associated with longer half-life (KW *p* < 10^-53^, **Fig. 3b**). Surprisingly, promiscuous oligomerization as attached particles have a mild effect in half-life elongation (KW *p* > 0.05). We perform two additional analyses to confirm this notion. First, we compare the half-life distributions of monomeric, central and attached proteins and observe significant differences in a manner 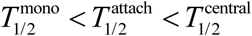, at similar ranges of *ACO* (**Fig. 3c**). Second, we classify the 2487 oligomeric proteins into three groups: proteins that participate in ≥1 complexes as central particles only (g1), those that contribute to ≥1 complexes as central and to ≥1 complexes as attached particles (g3) and those that participate in ≥1 complexes as attached particles only (g3) (**Fig. 3d-e**). Distributions of half-life differ significantly across these three groups in a manner 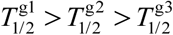 (KW *p* < 10^-41^, **Fig. 3f**). These results depict a proteome-wide tendency that central particles accomplish higher cooperative stability than attached particles upon complex formation.

### Cooperative stability of central subunits refers to burial of their short IDRs

Why do central particles achieve higher cooperative stability than attached particles upon oligomerization? We first check if this is because central particles exhibit higher *ACO* than attached particles. Notably, *ACO* distributions across g1, g2 and g3 do not differ significantly (KW *p* = 0.09 **, Fig. 4a**). It is already known that presence of sufficiently long terminal (∼30 residues) and internal (∼40 residues) IDRs, that can engage with proteasome, is associated with significantly shorter half-lives^12^. Interestingly, a comparison of IDRs across g1, g2 and g3 reveals three key aspects. (i) Lengths of both terminal and internal IDRs (*L*__IDR__) differ significantly across the three groups in a manner 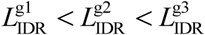 (KW *p* < 10^-3^, **Fig. 4b-d**). (ii) Central and attached proteins tend to have terminal IDRs shorter and longer, respectively, than the cutoff required for direct proteasomal engagement; (iii) both central and attached proteins exhibit internal disordered regions susceptible to direct proteasomal engagement (**Fig. 4b-d**). Since IDRs often get buried upon complex formation^26^, for crystal structures of 229 oligomeric proteins, we compare the percent of buried residues at terminal and internal IDRs (*B*__IDR__) upon oligomerization across the three groups. We find a statistically significant trend 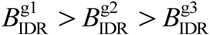 for internal IDRs only (KW *p* < 0.01, **Fig. 4e**).

**Figure 4.**
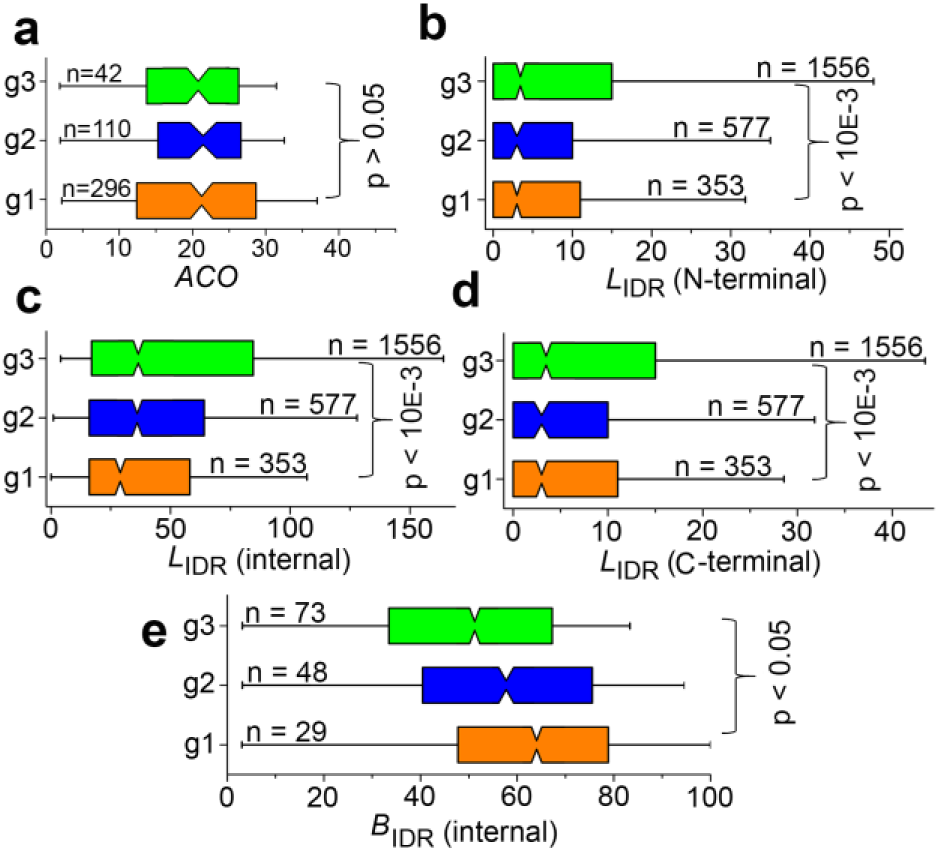
Differential cooperative stabilities of core and attached proteins. (**a**) Comparing *ACO* distributions of proteins across g1, g2 an g3 groups using permutation Kruskal-Wallis test. The three distributions do not differ significantly. Comparing the lengths of (**b**) N-terminal, (**c**) internal and (**d**) C-terminal intrinsically disordered regions of proteins across g1, g2 an g3 groups using permutation Kruskal-Wallis test. The length cutoff for terminal and internal disordered regions for direct proteasomal engagement is 30 and 40 amino acids (ref. 12). (**e**) Comparing the percent of internal disordered region burial proteins across g1, g2 an g3 groups using permutation Kruskal-Wallis test.

These two results suggest that higher cooperative stability of central subunits refer to their (i) significantly shorter terminal IDRs and higher burial tendency of internal IDRs upon complex formation, compared to those of attached particles. These attributes make central particles more likely candidates of escaping proteasomal engagement in the complexed state, compared to attached particles. Association with multiple complexes as central particles, is likely to elevate this probability of escaping degradation, explaining why promiscuous oligomerization leads to longer half-lives of central subunits. Cooperative stability thus acts a versatile and generic biophysical constraint to maintain oligomeric protein half-life (therefore abundance) according to their requirement in cellular machines.

### Complex self-assembly involve subunits with shorter half-lives binding late in the temporal order

The constituent subunits of macromolecular complexes follow evolutionarily conserved self-assembly pathways to organize themselves into complex functional machines^27–29^. A temporal order of subunit binding dictates that proteins binding early in the assembly order, remain in oligomeric state longer than those that bind late. Depending on the size of the complex, availability of subunits and cofactors, and efficiency of structural rearrangements to escape kinetic traps, self-assembly processes can continue from microseconds to several minutes^30,31^. This suggests that at least for large complexes, the temporal delay of subunit association can be long enough for proteasomal degradation rates to matter, resulting shorter subunit half-lives downward the assembly hierarchy. To test this, by an extensive literature search we collect the assembly hierarchy of 31 yeast complexes (Data S3) indispensable to central cellular processes such as replication, transcription, translation, cell cycle and transport. For distinct stages of subunit binding, we average the half-lives of respective subunits to estimate Spearman rank correlation (*rc*) with the temporal order. Consistent with our hypothesis, for 17 complexes with ≥3 stages of subunit binding (including ribosome, DNA and RNA Polymerases, kinetochore) we find *rc* = −1 (*p* < 0.05, **Fig. 5a**). For 18 additional complexes with only 2 stages of subunit binding (including nucleosome, DNA repair complex, mRNA decapping complex, Mitotic Checkpoint complex) half-life distributions also follow the same trend. 20S core particle of proteasome is the only exception, and being exception probably refers to significantly lower (MWU*p* < 0.05) *ACO* of α-subunits (21.8) that assemble prior to β-subunits (26.2).

**Figure 5.**
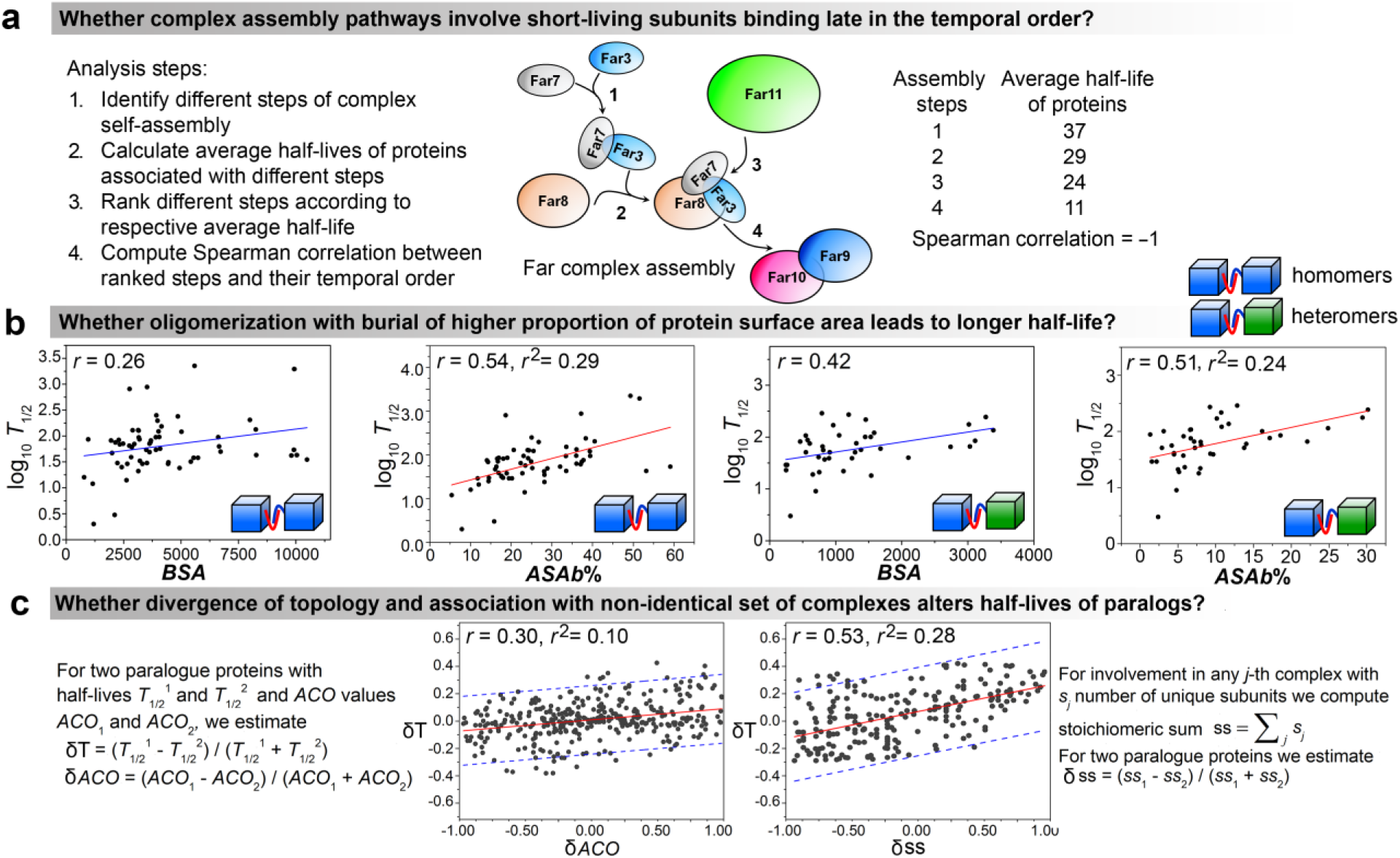
Assembly hierarchy and subunit buried surface area as regulators of protein half-life. (**a**) The computational pipeline to investigate whether and how assembly pathways of macromolecular complexes influence half-lives of different subunits binding in different temporal order. (**b**) Buried surface area (*BSA*) and percent of accessible surface area buried (%*ASAb*) of homomeric and heterodimeric complex subunits are correlated with subunit half-lives. Solid lines signify linear regressions in each case. (**c**) To investigate how structural (differential *ACO*) and functional divergence (association with non-identical sets of complexes), we compute three parameters shown in the figure: δT, δ*ACO* and δss. We find the linear regressions between δT and δ*ACO* and that between δT, δss. Solid lines signify linear regressions, dotted blue lines represent 95% prediction bands.

### Oligomerization with burial of higher proportion of protein surface area leads to longer half-life

Earlier studies on nonredundant heteromeric complexes with experimental binding kinetics data depicted positive and negative correlations of buried surface area (*BSA*) with association^32^ and dissociation^33^ rates. In these studies, a 500 Å^2^ to 3500 Å^2^ increment of *BSA* caused 10^10^-fold elevation of dissociation constant, that is expected to elevate the mean lifetime of a complex from milliseconds to hours^34^. These results suggest that for small dimeric complexes, higher *BSA* of the two subunits may lead to longer half-lives of the individual subunits. In other words, a positive correlation between *BSA* and *T*1/ 2 can be expected. Estimating the *BSA* from crystal structure data (Online Methods), we indeed find this positive correlation for homo and heterodimers (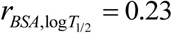, *p* < 0.01, **Fig. 5b**). This relationship applies to homomers up to dodecamers but not to heteromers any larger than dimers, probably because the temporal order of homomer dissociation largely follows the decreasing order of *BSA*^28^, which is not necessarily true for heteromeric complexes^29^. Use of percent of accessible surface area (*ASA*) buried (%*ASAb* = *BSA*×100 *ASA*) instead of *BSA*, elevates this correlationto 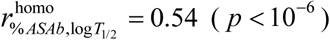 homo- and 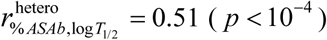 for heterodimers (**Fig. 5b**).

### Divergence of topology and of oligomerization promiscuity alters protein half-life in evolution

Structural determinants of protein half-life that we have analyzed so far are irrespective of the architecture of degradation machinery present in the cellular environment, which raises the question whether such attributes are exploited to alter protein half-life during evolution. Paralog protein pairs^12^ (arose from gene duplication) provide an excellent platform for such comparison between evolutionarily related proteins evolving under similar conditions. We observe a surprising result that divergence of native topology following gene duplication leads to altered half-lives of paralog pairs (**Fig. 5c**).

Gene duplication is often associated with loss and emergence of novel functions^35,36^. We identify a molecular signature of such functional diversification for 721 out of 1632 pre-identified yeast paralog pairs^12^ in terms of their oligomerization with non-identical sets (overlapping/nonoverlapping) of macromolecular complexes, which again, efficiently captures altered half-lives of these paralog pairs (**Fig. 5c**). Oligomerization with non-identical sets of complexes is associated with average ∼194 min variation of half-life, which is substantial, given ∼140 min average yeast doubling time during exponential growth^37^. Thus, altered half-life due to oligomerization with non-identical sets of macromolecular complexes could have a significant impact on the duration for which a protein can impart its function and thus affect cellular behavior.

## Discussion

How does the intrinsic structural features affect the lifetime of a protein? For over a decade this question has been of outstanding interest in molecular biology. The mechanistic details of proteasomal function led to the recognition of two factors to influence protein lifetime *in vivo*. Those include the presence of structural motifs promoting ubiquitinoylation^38^ and the presence of IDRs of sufficient size amenable to proteasomal engagement^12^. Our work extends the realm of these intrinsic structural features by distinguishing native topology of biological proteins and their potential to oligomerize into multicomponent complexes as master regulators of protein half-life in the cell. It is remarkable how simple geometrical considerations and oligomerization information appear to explain much of the differences in protein half-life over an entire genome, that includes nearly a thousand-fold variation of half-lives, and an enormous diversity of sequence, structure and function of the proteins compared.

The topological complexity of the protein fold (represented by *ACO*) plays a crucial role in determining the kinetics of protein folding^39^. We represent the first quantitative sketch of how the same factor acts as a molecular marker of their mechanical resistance and thereby captures variations of protein half-life. Further analysis shows that although overall degree of disorder of globular proteins also regulates their mechanical resistance, *ACO* plays the major deterministic role (**Text S1**). The role of long-range contacts in determining protein’s mechanical resistance was first revealed by comparing the unfolding patterns of ubiquitin and protein L, those feature similar fold class, but the former requires ∼70 pN higher force to unwind^21^. The terminal segments, by which the two proteins were pulled, make similar number of contacts with the hydrophobic core of both proteins, but the number of long-range contacts made between terminal regions of protein L and its hydrophobic core are significantly fewer than those for ubiquitin. This suggested that protein’s mechanical stability emerges from how the terminal—that is being pulled— is globally and cooperatively stabilized across the structure^21^. We generalize this concept in terms of a surprising correlation between mechanical resistance and *ACO*, that is further informative to capture half-life variations of thousands of proteins across yeast genome. The relationship between *ACO* and mechanical resistance may be the missing link to rationalize a wide range of observations regarding force-induced protein remodeling. The observation that native state *ACO* of β-sheet proteins is significantly higher than that of α-helix proteins of similar lengths (**Text S1**), indicates why the latter is mechanically weaker than the former^40,41^. Variation of *ACO* in different domains of multidomain proteins reflects their mechanical anisotropy (require unequal forces to unwind), and in turn, their directional preference to proteasomal degradation^22^.

For oligomeric proteins, sequestration into multimeric complexes itself warrants escaping proteasomal degradation to some extent^17–20^, resulting much weaker correlations between half-life and native topology. This correlation is even more weaker for proteins that remain disordered in monomeric state (**Text S1**). The role of cooperative stability to elongate protein lifetime in the complexed state was extrapolated multiple times in the past^17–20^, but this notion receives its first genome-scale assessment only in this study. Results depict that the impact of cooperative stability is generic but versatile. The generic nature is likely inherent to the fact that sequestration into complexes buries the disordered segments amenable to proteasomal engagement^26^, making oligomers more likely candidates of escaping proteasomal degradation compared to monomers having similar *ACO*. And the versatility is likely achieved by varying the temporal window of proteins being in the oligomeric state. This is attained by at least four mechanisms, (i) promiscuity of oligomerization (elevates the probability of finding the protein in oligomeric state), (ii) pervasive or temporary attachment with the complexes (central particles are permanent members of the complex and have longer half-lives than attached particles that are temporary members), (iii) temporal order of binding in the self-assembly pathway (early binding proteins remain in oligomeric state longer than late binding proteins) and (v) surface area buried upon binding (larger surface area ensures slower dissociation and hence longer half-life). This versatility of cooperative stability is believed to be important for the robustness and evolvability of genetic circuits^20^. These results further suggest that complex lifetime should have a strong influence on half-lives of its constituent subunits, challenging protein biochemists to assess this concept to direct experimental testing.

We observe that structural divergence upon gene duplication and association with differential set of macromolecular complexes influence half-lives of paralogue protein pairs. It also suggests a mechanism for divergence of half-life among orthologous proteins between species. Earlier, evolutionary variations leading to alteration of disordered regions was suggested to provide a simple evolutionary mechanism for fine-tuning protein lifetime according to regulatory sub-functionalization of paralogous proteins^12^. Our results suggest that fine-tuning protein half-life can also be achieved by harboring genetic variants that encode proteins with altered structural geometry compared to the wild-types^42^. Such evolutionary innovations are believed to manipulate regulatory schemes in genetic circuits to foster evolvability^43^.

In summary, our results reflect a complex interplay among versatile biophysical constraints associated with native topology, assembly, and oligomerization of biological macromolecules maintaining protein half-life in the cell. Native topology and oligomerization of proteins into multimeric complexes are independent of the architecture of the degradation machinery, and therefore, these factors are expected to be in effect equivalently in all living organisms.

## Acknowledgements

The authors sincerely acknowledge Vladimir Uversky (University of South Florida) for his critical reading and many useful suggestions. During the analysis phase of the work, we also acknowledge the constructive discussions with Tanaya Ray (Harischandra Research Institute).

## Author contributions

S.M. and S.K. conceptualized and designed research, S.M. collected and curated data and performed all the analyses, S.M., and S.K. discussed and interpreted the results and wrote the manuscript.

## Supplementary data

Protein Mechanical Resistance Data used in this study: Data S1, Yeast Complexome: Data S2, Complex assembly pathway database: Data S3, Paralogue Data: Data S4

## Online Methods

### Protein half-Life Data

Here we use the filtered *in vivo* protein half-life data for *Saccharomyces cerevisiae*, earlier analyzed by Madan Babu and co-workers^12^. This dataset includes the half-lives of 3273 yeast proteins, originally measured by Belle et al.^1^. Belle et al.^1^ measured protein half-lives by first inhibiting protein synthesis in exponentially growing yeast cells with the antibiotic cycloheximide and then monitoring the abundance of each C-terminally TAP-tagged protein in the yeast genome by quantitative Western blotting at three different time points.

### Protein structure data

On 19^th^ April 2017, we downloaded 2853 yeast protein X-ray crystallographic structures from Protein Data Bank^44^. Structures included in this dataset comprise only the annotated *Saccharomyces cerevisiae* proteins, obtained from the *Saccharomyces* Genome Database^45^. This initial dataset is further filtered based on two criteria: (i) the X-ray resolution is ≤ 3.0 Å and (ii) at least 75% of the primary chain is present in the electron density map, (iii) for multiple structures of the same protein, satisfying both the above criteria, we chose the highest-resolution structure. This filtering leaves us with only 267 crystal structures with ≤ 3.0 Å resolution. This dataset is too small, compared to the proteome-level half-life data of 3273 yeast proteins. Hence a reliable proteome-wide tendency cannot be expected to be derived by analyzing these 267 proteins only. Therefore, we also downloaded 5847 modeled yeast proteins from ModBase^46^. ModBase is a database of comparative protein structure models, calculated by a standardized automated comparative protein structure modeling pipeline^46^. In this pipeline, a structure model of the protein of interest in build based on one or more template structures having a certain degree of sequence identity with the protein of interest. A model is considered to be reliable (have a reliable fold assignment) if it is evaluated within the following thresholds by at least one of these model evaluation criteria^46^: (i) MPQS (ModPipe Quality Score) ≥ 1.1, (ii) TSVMod NO35 (estimated native overlap at 3.5 Å) ≥ 40%, (iii) GA341 (concerns the correct 3D coordinate assignments of the Cα atoms) ≥ 0.7, (iv) E-value (significance of the alignment between the target and the template by PSI-BLAST^47^) < 0.0001, and (v) zDOPE < 0 (for understanding the theoretical development of these parameters, please refer to ref. 46). We include a model structure in our structure dataset based on the following criteria: (i) the modeled region covers ≥ 75% of the protein length, (ii) MPQS (ModPipe Quality Score) ≥ 1.1, (iii) TSVMod NO35 (estimated native overlap at 3.5 Å) ≥ 40%, (iv) GA341 ≥ 0.9, (v) PSI-BLAST E-value between model and template structures is < 10^-8^, and (vi) zDOPE < 0. After applying these constraints, we are left with reliable model structures of 1003 proteins.

### Protein intrinsic disorder data

We have used the intrinsic disorder data of 3273 yeast proteins (those with available half-life data), earlier predicted by Madan Babu and co-workers^12^. The authors used three complementary methods^48–50^ for inferring residue-level disorder tendency of each yeast protein.

### Protein mechanical unfolding data

The mechanical unfolding of globular proteins upon pulling the amino acid chain (similar to that occurs upon proteasome engagement) has been addressed by Atomic Force Microscopy experiments and by computer simulations. We have used three datasets in our work (Data S1). The first group (G1) includes 16 proteins unfolded in Atomic Force Microscopy experiments, by pulling the polypeptide chains at 600 nm/s (11 proteins) and 300 nm/s (5 proteins) speeds. This dataset is collected from Brockwell et al.^50^ and Sulkowska and Cieplak^51^. The second group (G2) includes 27 proteins unfolded in all-atom computer simulations. Eleven proteins (G2A) were pulled from the N-terminal at 5 ×10^7^ nm/s speed (C-terminal fixed). This data is also collected from Sulkowska and Cieplak^51^. Sixteen proteins (G2B) were pulled from N- and C-terminal separately at 5 ×10^6^ nm/s speed, keeping the other terminal free, thus allowing the substrate to rotate and adopt a less obstructive orientation for unfolding (as happens during degradation). This data is collected from the work of Wojciechowski et al.^52^.

In addition, we have collected experimental mechanical unfolding data for three proteins, *Dictyostelium discoideum* filamin^53^, yellow and green fluorescent proteins^54^ and Ig27 domain of titin^55^, each depicting alterations of mechanical resistance upon point-mutations in the native protein. This data is used to verify whether enhancement/reduction of mechanical resistance in these cases are associated with respective increase/decrease of contact order.

### Absolute Contact Order estimation (*ACO*)

The absolute contact order (*ACO*) of a protein structure is defined as the average amino acid separation of 3D contacts^56^:

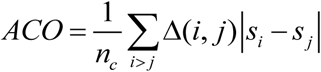

where *n*_*c*_ is the total number of residue-residue contacts, *s*_*i*_ and *s* _*j*_ are the sequence positions of residues *i* and *j*, and Δ(*i*, *j*) is the selection criteria that includes *i* and *j* into analysis only if they are in contact and if |*i* - *j*| ≥ 4. This |*i* - *j*| ≥4 criterion ensures that contacts included in *ACO* estimation reflect 3D topology of the proteins, rather than secondary structures. We defined a residue contact between a pair of residues when the distance between any two atoms from the residue pair is less than the sum of their van der Waals radii plus 0.5 Å cut-off distance^57^.

### Accessible and buried surface area calculation

The Surface Racer program^58^ is used to calculate the solvent accessible and buried surface areas of the proteins, with probe radius taken to be 1.4 Å, which resembles the radius of one water molecule. We calculated the solvent accessible surface area (*ASA*) of the two interacting partners separately (in their complexed conformation) and in associated state. If the *ASA* of the two partners are *A*1 and *A*2 and of their associated structure is *A*3, then buried surface area (*BSA*) is defined as (*A*1+*A*2-*A*3)/2.

### Proteome-wide screening for macromolecular complexes

For a proteome-wide screening of yeast macromolecular complexes, we begin with downloading the previously published dataset of 491 yeast complexes by Gavin et al.^24^ and 412 complexes included in the Complex Portal of the IntAct Molecular Interaction Database^25^. The database presented by Gavin et al.^24^ was the first proteome-wide screening for macromolecular machines in yeast, using tandem-affinity-purification method coupled to mass spectrometry (TAP–MS) to all 6,466 ORFs of *Saccharomyces cerevisiae*. Entries in Complex Portal^25^ are based on manual curation of widespread experimental data depicting direct physical association between complex subunits, such as affinity chromatography, chromatin immunoprecipitation, coimmunoprecipitation, two hybrid fragment pooling, tandem affinity purification, electron microscopy and x-ray crystallography. We (i) carefully compare the entries in these two databases, (ii) further curate the information therein (regarding the existence of the complex and its subunit composition) based on extensive protein-by-protein literature search of published experimental data and (iii) add new complexes in the set (along with subunit composition information) accordingly. We particularly look for reports concerning (i) different isoforms of a given complex and (ii) its temporary attached particles, such as chaperons and assembly co-factors. Homology-based predicted complexes are disregarded and only experimentally verified complexes are considered. We finally identify 80 homomeric and 805 heteromeric complexes. **Data S2** includes the association information of different proteins with different macromolecular complexes along with literature reference.

Using this data, first for any given complex, we classify the subunits into two groups: (i) functional subunits present in the matured complex (if different isoforms exist they are present in most of the isoforms) are called central, the remaining (ii) temporary attached proteins such as assembly cofactors and subunits present only in some of the isoforms of a complex are called attached particles.

In addition to subunit composition, we further look for literature evidence concerning self-assembly of macromolecular complexes. By an extensive literature search we collect the assembly hierarchy of 35 yeast complexes (Data S3) indispensable to central cellular processes such as replication, transcription, translation, cell cycle and transport.

### Yeast paralogue data

Yeast paralog pairs were obtained from the work of van der Lee et al.^12^; authors generated the paralogue set by an all-against-all sequence comparison using BLASTClust^47^. They added more divergent paralogs from the yeast whole-genome duplication event^61^.

### Statistical Analysis

All the statistical analyses are performed using in-house Python scripts and PAST software package^62^.

## Data S1

### Group G1

Dataset of proteins unfolded in Atomic Force Microscopy experiments. The Protein Data Bank (PDB) code is listed if the corresponding structure of the protein is available in PDB. *F*_max_ represent the peak unfolding force required to unwind the corresponding protein, *v*_p_ is the chain pulling speed. Literature reference for each protein is provided.

| Protein | PDB | $F_{\max}$ (pN) | $v_p$ (nm/s) | Reference |
| --- | --- | --- | --- | --- |
| α-spectrin | 1U4Q | 30 | 300 | 1, 2 |
| α-spectrin R16 | 1AJ3 | 54 | 300 | 3 |
| α-spectrin13-18, 18-21 | 1U4Q | 26 | 300 | 1, 2, 4 |
| β-spectrin1-4 | 1S35 | 27 | 300 | 1, 2, 5 |
| α-actin1-4 | 1HCI | 38 | 300 | 2, 4 |
| lipoyl domain of aceF | 2K7V | 15 | 600 | 6 |
| C2A | 1DQV | 60 | 600 | 7 |
| T4 lysozyme | 1B6I | 64 | 600 | 8 |
| <sup>10</sup>FNIII | 1FNF | 74 | 600 | 9, 10 |
| Calmodulin | 1CFC | 80 | 600 | 7 |
| <sup>13</sup>FNIII<sub>27</sub> | 1FNH | 89 | 600 | 10 |
| TNFn | 1TEN | 113 | 600 | 11 |
| <sup>1</sup>FNIII<sub>27</sub> | 1OWW | 120 | 600 | 10 |
| <sup>12</sup>FNIII<sub>13FNIII</sub> | 1FNH | 124 | 600 | 10 |
| I27 | 1TIT | 127 | 600 | 12 |
| FNIII | 2N1K | 220 | 600 | 10 |

### Group G2A

Dataset of proteins unfolded in Computer simulations by pulling the N-terminal, while keeping the other terminal fixed. The Protein Data Bank (PDB) code is listed if the corresponding structure of the protein is available in PDB. *F*_max_ represent the peak unfolding force required to unwind the corresponding protein, *v*_p_ is the chain pulling speed. Literature reference for each protein is provided.

| Protein | PDB | $F_{\max}$ (pN) | $v_p$ (nm/s) | Reference |
| --- | --- | --- | --- | --- |
| I1 oxidized | 1GCG | 2397 | $5 \times 10^7$ | 1 |
| I1 reduced | 1GCG | 2090 | $5 \times 10^7$ | 1 |
| I27 | 1TIT | 2479 | $5 \times 10^7$ | 2 |
| I27 | 1TIT | 2040 | $5 \times 10^7$ | 2, 3 |
| <sup>7</sup>FNIII | 1FNF | 1638 | $5 \times 10^7$ | 4, 5 |
| <sup>10</sup>FNIII | 1FNF | 1580 | $5 \times 10^7$ | 4, 5, 6 |
| Bovine | 1V9E | 3000 | $5 \times 10^7$ | 7 |
| Cad1 | 1EDH | 1850 | $5 \times 10^7$ | 5 |
| Cad2 | 1EDH | 1970 | $5 \times 10^7$ | 5 |
| Cell adhesion VCAM1 | 1VSC | 2050 | $5 \times 10^7$ | 5 |
| Cell adhesion VCAM2 | 1VSC | 1620 | $5 \times 10^7$ | 5 |

### Group G2B

Dataset of proteins unfolded in Computer simulations by pulling N- and C-terminal separately, while keeping the other terminal free. The Protein Data Bank (PDB) code is listed if the corresponding structure of the protein is available in PDB. *F*_max_ represent the peak unfolding force required to unwind the corresponding protein, *v*_p_ is the chain pulling speed. Literature reference for each protein is provided.

| Protein | PDB | $F_{\max}$ (N-terminal)<br>(pN) | $F_{\max}$ (C-terminal)<br>(pN) | $v_p$ (nm/s) | Reference |
| --- | --- | --- | --- | --- | --- |
| Cohesin | 1AOH | 495 | 407 | $5 \times 10^6$ | 1 |
| Green fluorescent protein | 1GFL | 253 | 253 | $5 \times 10^6$ | 1 |
| Nudix | 1VCD | 440 | 385 | $5 \times 10^6$ | 1 |
| Cytolysin | 1O72 | 286 | 341 | $5 \times 10^6$ | 1 |
| Cytolysin | 1GWY | 308 | 341 | $5 \times 10^6$ | 1 |
| Purine nucleoside phosphorylase | 1ODI | 352 | 374 | $5 \times 10^6$ | 1 |
| Lectin | 1Y2X | 319 | 286 | $5 \times 10^6$ | 1 |
| Lutheran glycoprotein | 2PF6 | 297 | 407 | $5 \times 10^6$ | 1 |
| Purine nucleosidase phosphorylase | 1OTX | 297 | 341 | $5 \times 10^6$ | 1 |
| ADP-ribose pyrophosphatase | 2DSD | 352 | 286 | $5 \times 10^6$ | 1 |
| Purine nucleoside phosphorylase | 1NW4 | 319 | 363 | $5 \times 10^6$ | 1 |
| Dihydrofolate reductase | 1U71 | 275 | 253 | $5 \times 10^6$ | 1 |
| I27 domain of titin | 1TIT | 264 | 286 | $5 \times 10^6$ | 1 |
| Ribonuclease H | 1RIL | 363 | 231 | $5 \times 10^6$ | 1 |
| Barnase | 1BNR | 209 | 176 | $5 \times 10^6$ | 1 |

### Proteins with experimental evidence concerning altered mechanical resistance upon point mutations

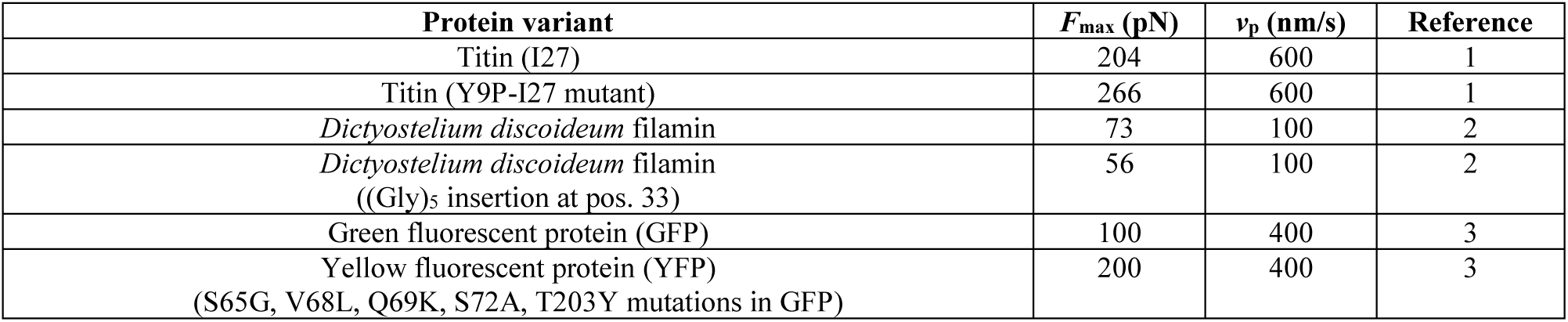

| Protein variant | $F_{\max}$ (pN) | $v_p$ (nm/s) | Reference |
| --- | --- | --- | --- |
| Titin (I27) | 204 | 600 | 1 |
| Titin (Y9P-I27 mutant) | 266 | 600 | 1 |
| *Dictyostelium discoideum* filamin | 73 | 100 | 2 |
| *Dictyostelium discoideum* filamin<br>((Gly)<sub>5</sub> insertion at pos. 33) | 56 | 100 | 2 |
| Green fluorescent protein (GFP) | 100 | 400 | 3 |
| Yellow fluorescent protein (YFP)<br>(S65G, V68L, Q69K, S72A, T203Y mutations in GFP) | 200 | 400 | 3 |

## Data S2 and Data S3

Available upon personal request to the authors.

## Text S1

**Protein’s mechanical resistance correlates with their fraction of disordered residues as well**

It is a well-established fact that presence of disordered regions is associated with weaker mechanical resistance of biological proteins and proteasomes exploit this attribute by preferring the disordered termini of substrate proteins as initiation sites of forced unwinding^1^. Since the percent of disordered residues (*PDR*) present within a protein can be considered as a measure of their overall stability^2,3^, we aim to find a quantitative sketch of how *PDR* correlates with protein’s mechanical resistance. We estimate the *PDR* of proteins included in G1, G2A and G2B datasets using DISOPRED3 algorithm^4^ and obtain statistically significant negative correlations with peak unfolding forces.

Linear regression for G1 group:

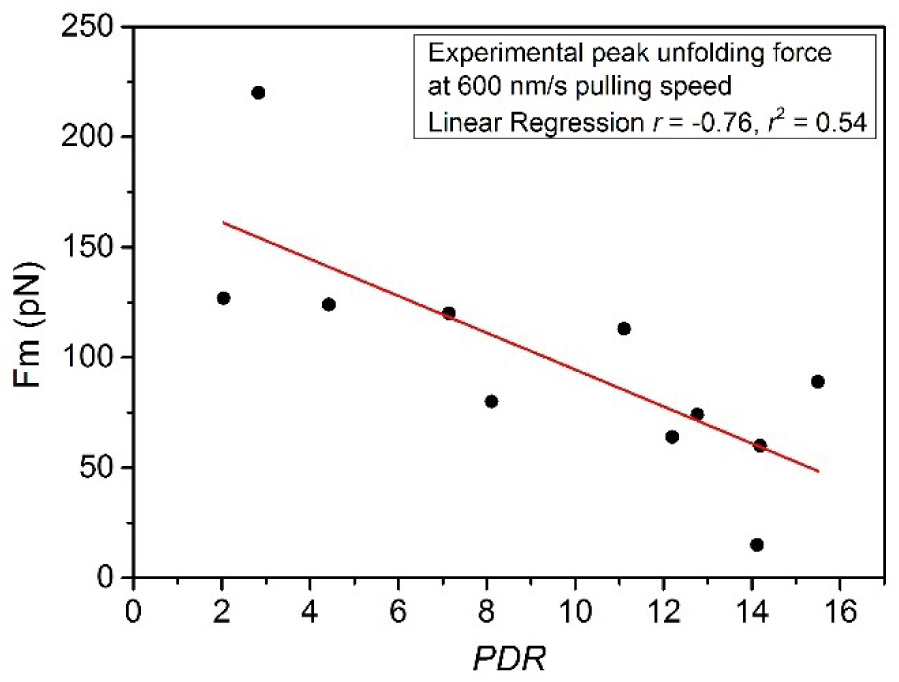

Linear regression for G2A group:

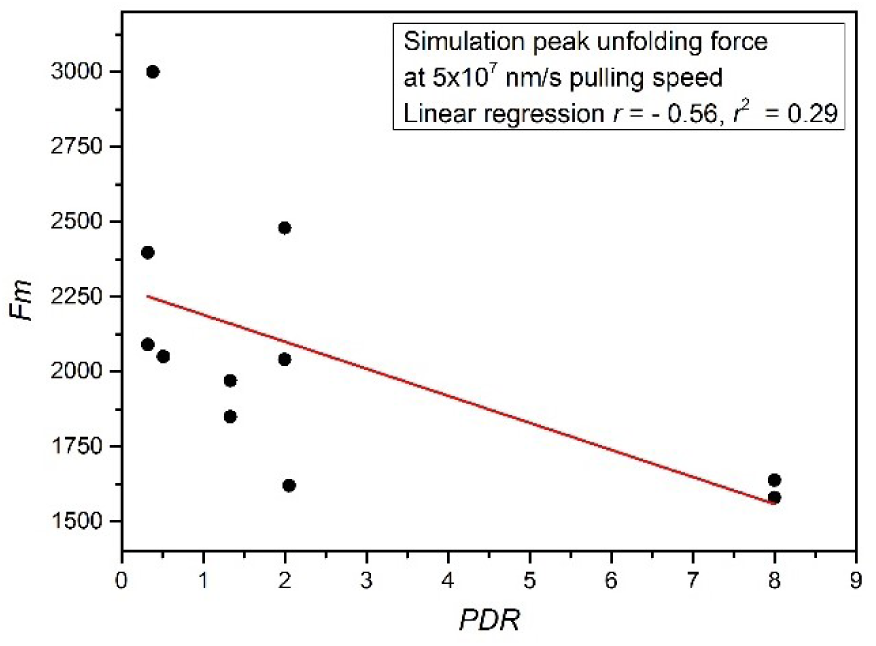

Linear regression for G2B group:

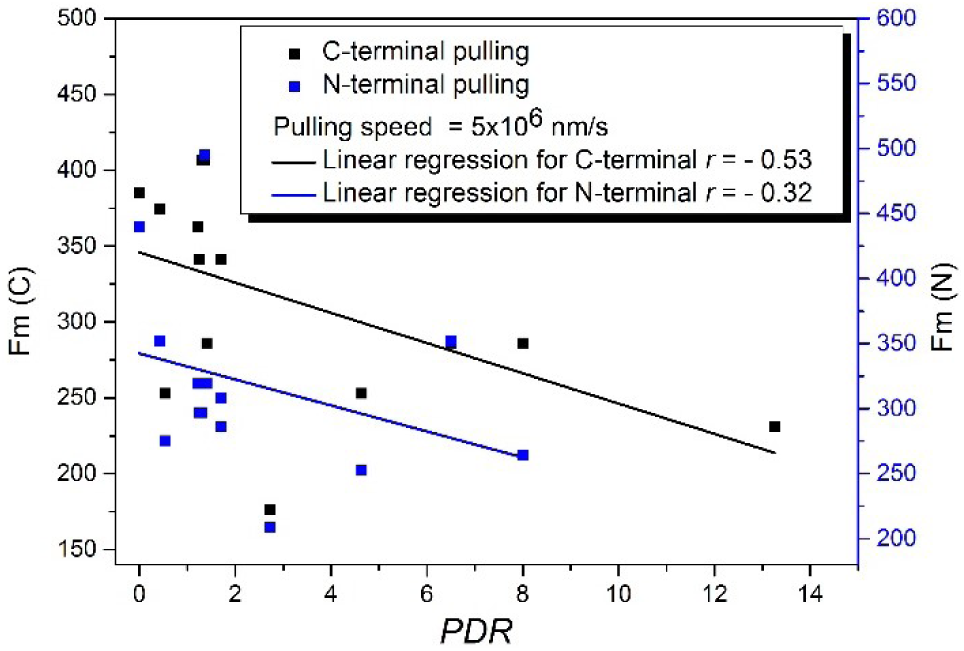

The correlation between *PDR* and mechanical resistance is basically quantitatively presents the already established concept that proteins with higher degree of disorderness would unfold easily. Though this issue has been repetitively tested and validated in multiple experiments, a simple mathematical sketch has been missing until now.

## *ACO*, not *PDR*, is the major determinant of protein’s mechanical resistance

In the main text, we have shown that higher *ACO* is associated with higher mechanical resistance of substrate proteins. Here we observe another striking fact that overall degree of disorderness, measured as *PDR*, contributes to protein’s mechanical resistance as well.

If *ACO* and disorder both determine mechanical resistance, what are their unique contributions (if the other is absent) to the correlation with mechanical resistance? We have calculated the partial correlations of *ACO* and *PDR* with mechanical resistance to answer this question. If both A and B correlate with C, partial correlation between A and C excludes the effect of B to estimate A’s unique contribution.

**Results for G1 set (* signifies correlation *p* value < 0.05)**

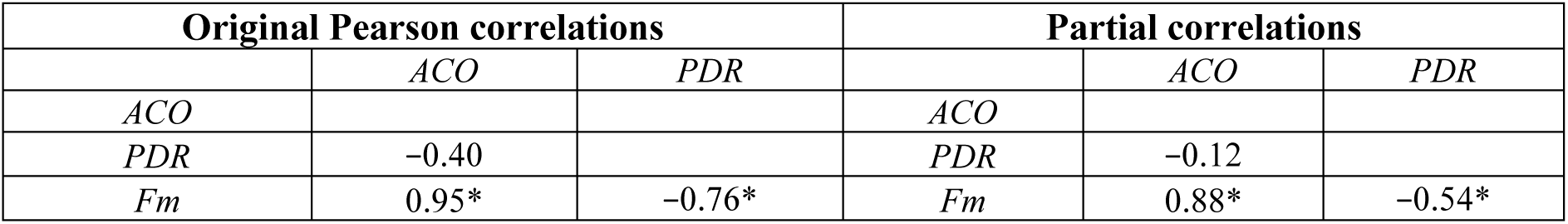

**Results for G2A set (* signifies correlation *p* value < 0.05)**

**Results for G2A set (\* signifies correlation *p* value < 0.05)**
| Original Pearson correlations |  |  | Partial correlations |  |  |
| --- | --- | --- | --- | --- | --- |
|  | *ACO* | *PDR* |  | *ACO* | *PDR* |
| *ACO* |  |  | *ACO* |  |  |
| *PDR* | -0.37 |  | *PDR* | -0.09 |  |
| *Fm* | 0.74* | -0.60* | *Fm* | 0.68* | -0.47 |

**Results for G2B set (* signifies correlation *p* value < 0.05)**

**Results for G2B set (\* signifies correlation $p$ value < 0.05)**
| Original Pearson correlations |  |  | Partial correlations |  |  |
| --- | --- | --- | --- | --- | --- |
|  | *ACO* | *PDR* |  | *ACO* | *PDR* |
| *ACO* |  |  | *ACO* |  |  |
| *PDR* | -0.53* |  | *PDR* | -0.18 |  |
| *Fm* | 0.89* | -0.52* | *Fm* | 0.85* | -0.21 |

The partial correlation between *ACO* and *PDR* vanishes when we exclude the effect of mechanical resistance. This proves the correlation between *ACO* and *PDR* is merely an indirect correlation. The fascinating observation is that, even after excluding the effect of *PDR*, there is only a little reduction of the correlation between mechanical resistance and *ACO*. Conversely, if we exclude the effect of *ACO*, the correlation between mechanical resistance and *PDR* drops severely. This clearly dictates that at least for small globular proteins *ACO* is the major deterministic factor of mechanical resistance, while *PDR* has a minor effect. This result may be a result of the fact that proteins included in these datasets are all globular proteins with a few disordered residues at their termini. Disorder may play much stronger roles in proteins with longer disordered segments.

### Complex subunits, that remain structured in monomeric state, depict stronger correlations between *ACO* and half-life

Gunasekaran et al.^5^ showed that a simple plot (Nussinov plot) of length-normalized Buried Surface Area (*BSA*/*L*) versus length-normalized Accessible Surface Area (*ASA*/*L*) of complex subunits (at complexed state) can tell us whether a subunit of interest remains unstructured or structured at monomeric state. We exploit this concept to infer the disorder/order status of complex subunits with resolved crystal structures at monomeric state (see Online Methods).

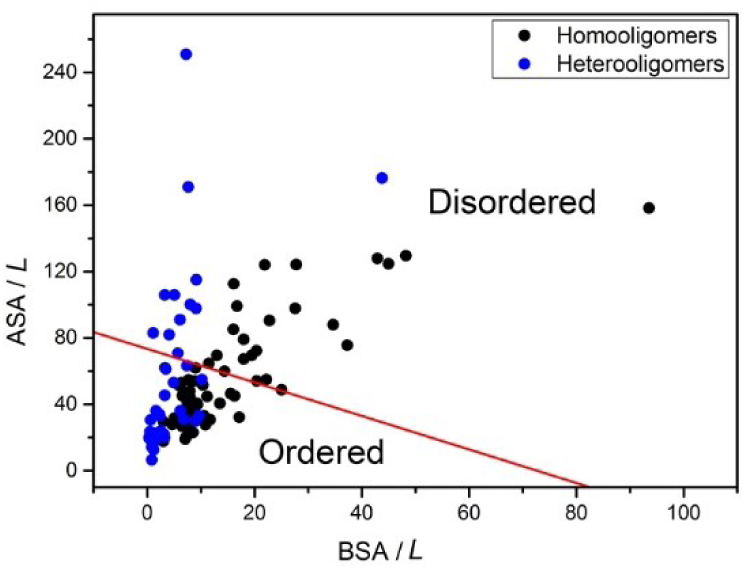

Using this plot, we estimate the correlations between half-life and absolute contact order (*ACO*) for subunits (i) those that remain unstructured and (ii) structured in monomeric state.

Linear regressions for homo- and heteromeric subunits that remain disordered in monomeric state:

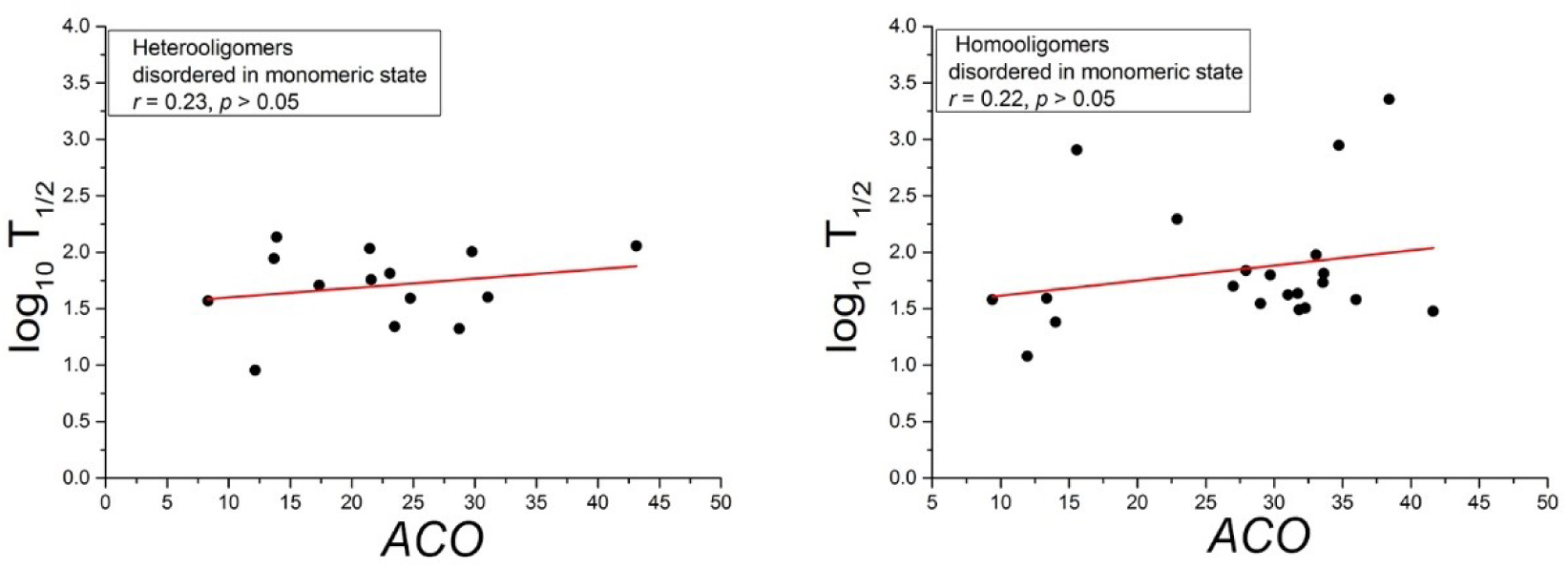

Linear regressions for homo- and heteromeric subunits that remain structured in monomeric state:

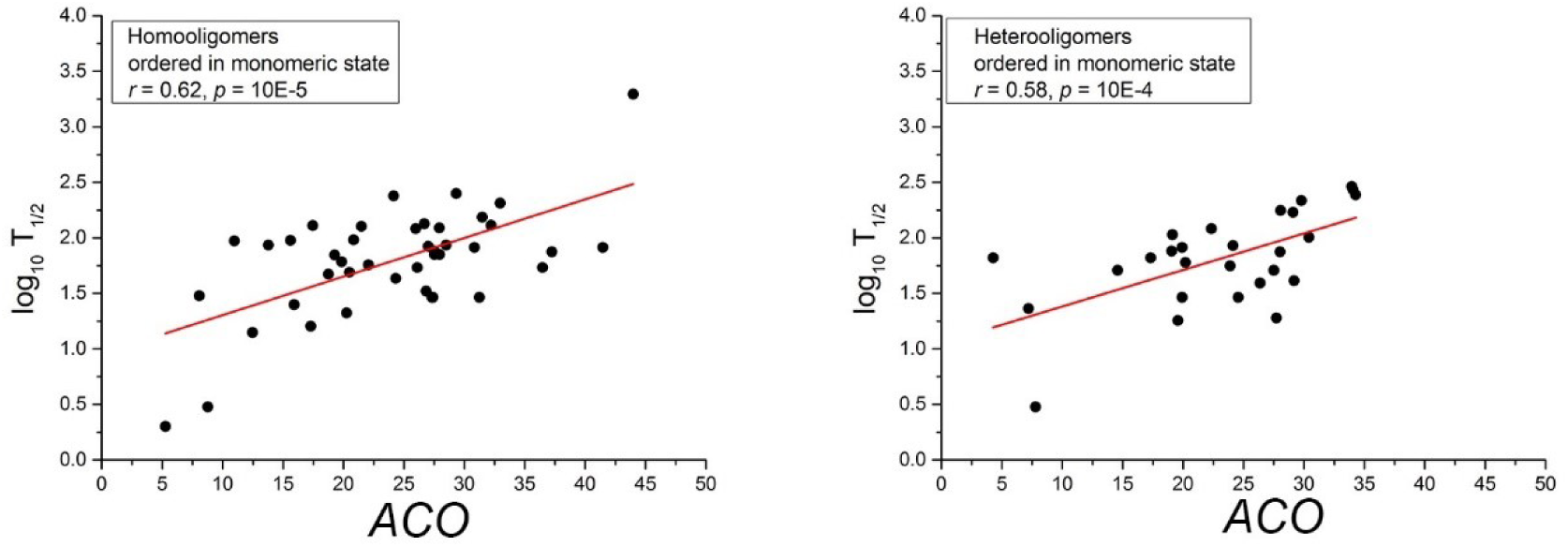

Nussinov plot leads us to some very interesting conclusions. First, *ACO* is a much stronger regulator of half-life for oligomeric proteins that remain structured in monomeric states as well, compared to those that remain disordered at monomeric state. It is a well-known fact that oligomeric proteins degrade much faster in monomeric state. So, for proteins that remain structured prior to degradation, *ACO* stands as a marker of mechanical resistance and thus affects half-life. For proteins, that become unstructured at monomeric state, one cannot expect a correlation between complexed state *ACO* and half-life.

### Stronger correlations between BSA and half-life is obtained for complex subunits that remain structured in monomeric state, compared to those that remain disordered

In the main text, we have shown that *BSA* acts as a marker of dissociation rate of complex subunits. Because oligomeric proteins degrade much faster in monomeric state, proteins that dissociate slowly from complexes, have longer half-life. We now ask whether this relationship depends on the fact that some oligomeric proteins remain structured and others remain disordered in monomeric state.

Linear regressions between *BSA* and half-life for homo- and heteromeric complex subunits that remain disordered in monomeric state:

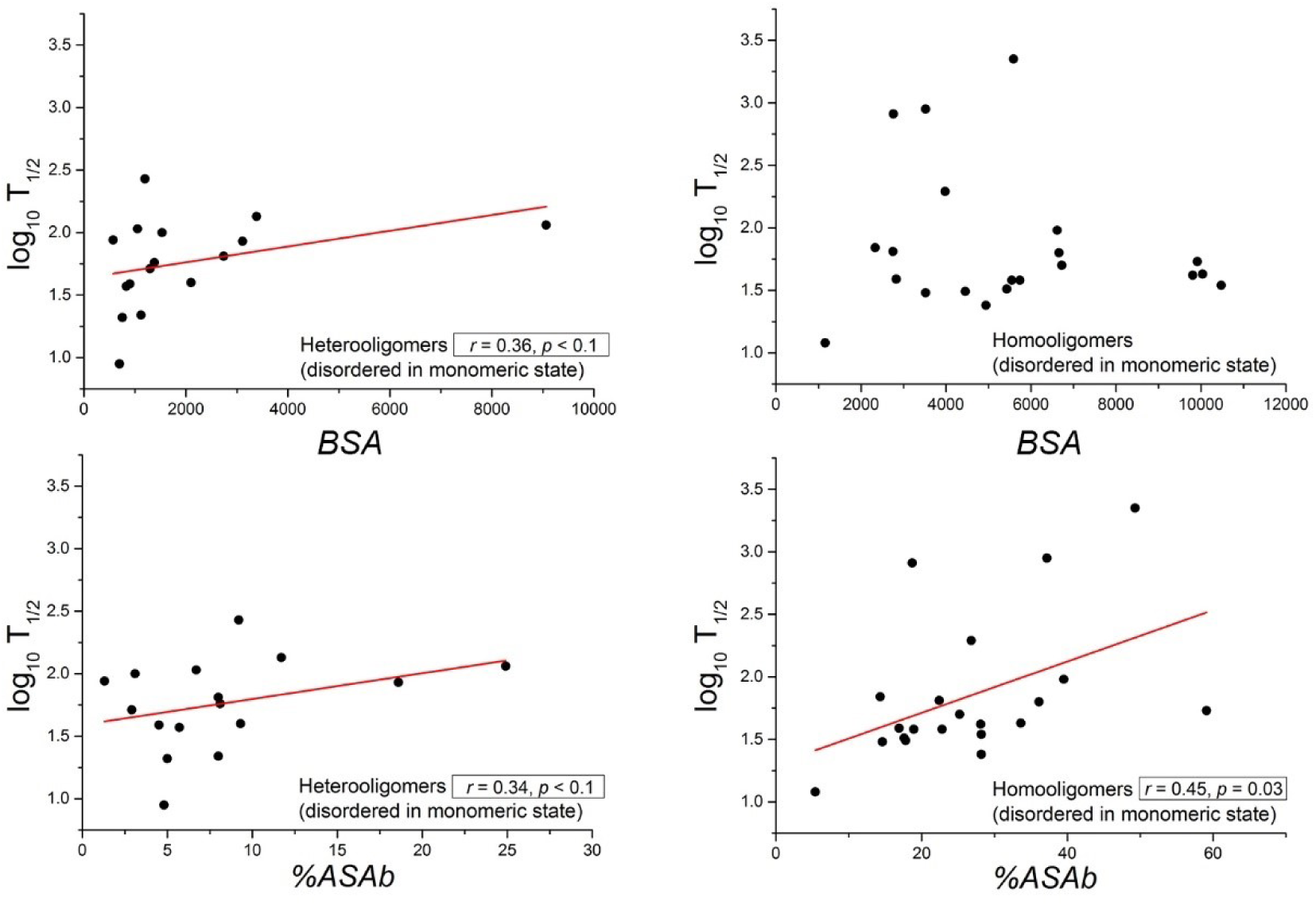

Linear regressions between *BSA* and half-life for homo- and heteromeric complex subunits that remain structured in monomeric state:

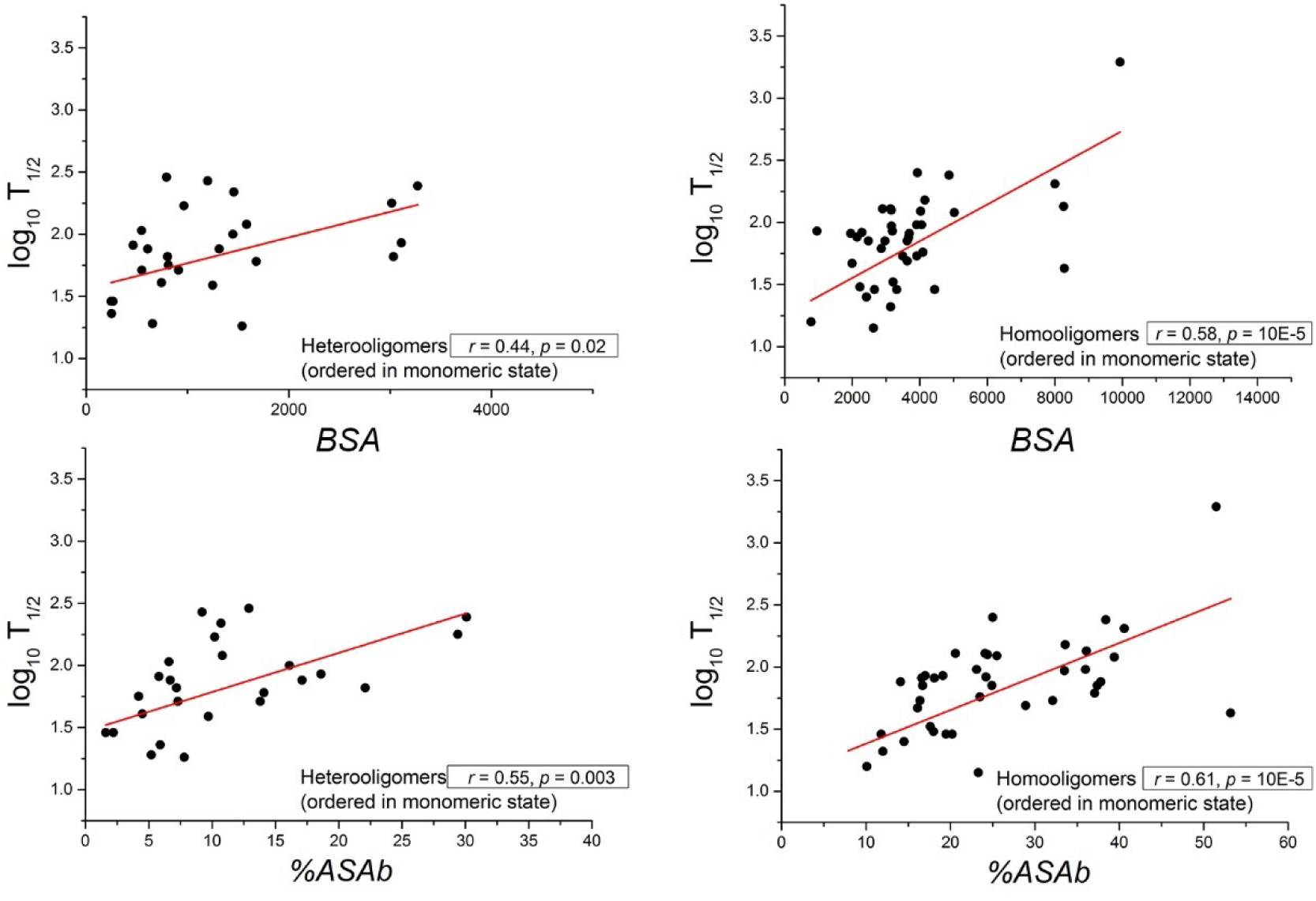

Nussinov plot again leads us to some very interesting conclusions. Proteins that obtain stable 3D structure only after binding depict weaker dependency with *BSA*. Just as we noticed in the main text, the correlation gets improved when we consider percent of ASA buried (%*ASAb*) instead of *BSA*. It has been previously shown that proteins with higher disorderness can even get degraded directly from complexes, if proteasomes can access their segments that remain disordered even in complexed state. This probability of proteasomal engagement is expected to be much weaker for proteins that are remain ordered even in monomeric state, explaining the *BSA* dependency.

